# Altering rRNA 2’O-methylation pattern during neuronal differentiation is regulated by FMRP

**DOI:** 10.1101/2024.07.25.605157

**Authors:** Michelle Ninochka D’Souza, Naveen Kumar Chandappa Gowda, Nivedita Hariharan, Syed Wasifa Qadri, Dasaradhi Palakodeti, Ravi S Muddashetty

**Author notes:** Equal contribution (Michelle Ninochka D’Souza, Naveen Kumar Chandappa Gowda).

## Abstract

The Fragile X Messenger Ribonucleoprotein (FMRP) is a selective RNA-binding protein that localizes to the cytoplasm and the nucleus. The loss of FMRP results in Fragile X Syndrome (FXS), an Autism Spectrum Disorder. FMRP interacts with ribosomes and regulates the translation of mRNAs essential for neuronal development and synaptic plasticity. However, the biochemical nature of this translation regulation is unknown. Here we report that a key feature of FMRP-mediated translation regulation during neuronal differentiation is modulating the 2’O-methylation of ribosomal RNA. 2’O-methylation, facilitated by C/D box snoRNAs in the nucleus, is a major epitranscriptome mark on rRNA, essential for ribosome assembly and function. We found that FMRP influences a distinct rRNA 2’O-Methylation pattern across neuronal differentiation. We show that in H9 ESCs, FMRP interacts with a selected set of C/D box snoRNA in the nucleus resulting in the generation of ribosomes with a distinct pattern of rRNA 2’O-Methylation. This epitranscriptome pattern on rRNA undergoes a significant change during the differentiation of ESCs to neuronal precursors and cortical neurons. ESCs display maximum hypomethylated residues on rRNA, which is eventually reduced in neuronal precursors and post-mitotic cortical neurons and this is correlated to the change in global protein synthesis among the states of differentiation. Importantly, this gradual change in the 2’O-methylation pattern during neuronal differentiation is altered in the absence of FMRP, which could impact neuronal development and contribute to dysregulated protein synthesis observed in Fragile X Syndrome. This also suggests the need for diversity in functional ribosomes during the early stages of development.

## Introduction

Dynamic change in the protein repertoire mediated by translation regulation has shown to be a critical determinant of Embryonic stem cell (ESCs) maintenance and differentiation^1,2^. Several factors including non-coding RNAs, RNA binding proteins, and epitranscriptomic modifications such as mRNA m6A modifications are shown to alter translation rates, thereby, regulating the protein repertoire of ESCs^3^. Recent work showed that rRNA modifications that change between cell types could potentially regulate translation by modulating the interaction between mRNA and the ribosomes ^4^. However, the role of epitranscriptomic modification at the rRNA level and its contribution to ESC differentiation are largely unexplored. 2’O-methylations are one of the major epitranscriptomic marks found on rRNA. In humans, C/D box small nucleolar RNAs (snoRNAs) guide the addition of 2’O-methylation on the ribose sugar of rRNA and this process can occur either co-transcriptionally or post-transcriptionally^5^. rRNA methylation is important for ribosome biogenesis as it helps in the folding of rRNA and assembly of ribosomes^6^. The idea of ribosomes being structurally and functionally uniform entities has been seriously challenged in recent years and the idea of ribosome heterogeneity is gaining wide acceptance^7–9^. Ribosome heterogeneity can be attributed to content and the modifications of both proteins and rRNA. Though the change in the protein composition of ribosomes was shown to have a regulatory role in translation, the consequence of rRNA-based ribosome heterogeneity based on translation regulation is largely unexplored^10^.

In our previous work, we showed that the differential pattern of rRNA 2’O-methylation in Shef4 hESCs was contributed by a large extent of hypomethylated residues. Further, our work also demonstrated that Fragile X Messenger Ribonucleo-binding protein (FMRP), an RNA binding protein, modulates this rRNA methylation at several specific sites generating a differential 2’O-methylation pattern^11^. Consequently, the differential 2’O-methylation pattern on the ribosome assists in the binding of FMRP, which plays an important role in the regulation of protein synthesis^11^. FMRP-mediated translation regulation is critical for brain development and functioning^12–14^. Consequently, the loss of FMRP results in a severe form of Autism Spectrum Disorder called Fragile X Syndrome, which is primarily characterized by intellectual disability^15^

In our current study, we investigated the changes and functional relevance of the 2’O-methylation pattern of rRNA in the maintenance and differentiation of hESCs to neural fates. Our results show that the translation rates are higher in NSCs compared to ESCs suggesting that cell state changes correlate with dynamic change in the rates of protein synthesis. We conclude that rRNA hypomethylation broadly correlates to lower translation as in the case of WT ESCs while rRNA hypermethylation correlates with increased translation as seen in WT NSCs. This correlation was also observed in FMR1 KO ESCs. Our study captures the significant change in the 2’O-methylation pattern during differentiation with a maximum number of hypo-methylated sites in the ESCs. Further, our results show changes in the 2’O-methylation of rRNA in translating and non-translating pools of the ribosome, suggesting that these methylation statuses might have a profound influence on translation rates. Our work also demonstrates that the changes in methylation patterns from ESCs to NSCs are particularly regulated by specific snoRNAs in association with FMRP. Together, this study provides insights into 2’O-methylation-dependent translation regulation mediated by FMRP and its importance in the differentiation of ESCs to neural lineages.

## Results

### The hypomethylated sites are maximum in ESC rRNA and significantly reduce as they differentiate into NSCs and neurons

We sought to investigate the changes in 2’O-methylation marks on ribosomal RNA during ESC differentiation to neuronal precursors and mature neurons. For this purpose, H9 ESCs were differentiated into Neural Stem Cells (NSCs) and finally into forebrain glutamatergic neurons through the inhibition of the SMAD signaling pathway (**Figure S1 A-C**). RNA from ESCs, NSCs, and differentiated neurons was subjected to RiboMethSequencing (RMS) to estimate the changes in the 2’O-methylation patterns across 18S and 28S rRNA as described previously^16^. In brief, 2 micrograms of total RNA were subjected to controlled alkaline hydrolysis followed by library preparation and sequencing (**Figure 1A**). The extent of 2’O-methylation of specific sites of the 18S and 28S rRNA are represented as methylation indices (MI). MI=1 indicates complete methylation at a particular site, while MI=0.1 indicates a methylation of only 10% of the rRNA population at a particular site. RiboMeth scores for ESCs, NSCs, and neurons indicate differential patterns of 2’O-methylation in 18S and 28S rRNA across these three stages of neural differentiation (**Figure 1B and 1C**). Our RiboMethSeq captured a total of 103 differentially methylated sites with 39 residues in 18S rRNA and 64 residues in 28S rRNA respectively (**Figure 1B and 1C**). Further, we captured two distinct patterns in our RMS score: a) Among the three differentiation stages, ESC rRNA displayed the highest number of sites with partial 2’O-methylation. This indicates that ESCs contain a maximum number of ribosomes having hypomethylated residues and b) the number of 2’O hypo-methylated residues decreases as the ESCs differentiate to NSCs and reduce even further as the NSCs undergo transition to post-mitotic neurons (**Figure 1B and 1C and Figure S1D and S1E**). Detailed information on sites that show a significant increase in methylation across ESC to neurons is provided in **Table 1**. Conversely, a few sites in 18S and 28S rRNA show a significant shift to hypomethylation in NSCs and neurons compared to ESC (indicated in **Table 2**), which is an opposite trend observed in the majority of the sites mentioned earlier. The pattern of 2’O-methylation obtained was distinct among H9 ESCs and the differentiated NSCs and neurons. However, the hypomethylated residues in H9 ESC were the same as those captured in our previous study using an alternate ESC line Shef4^11^

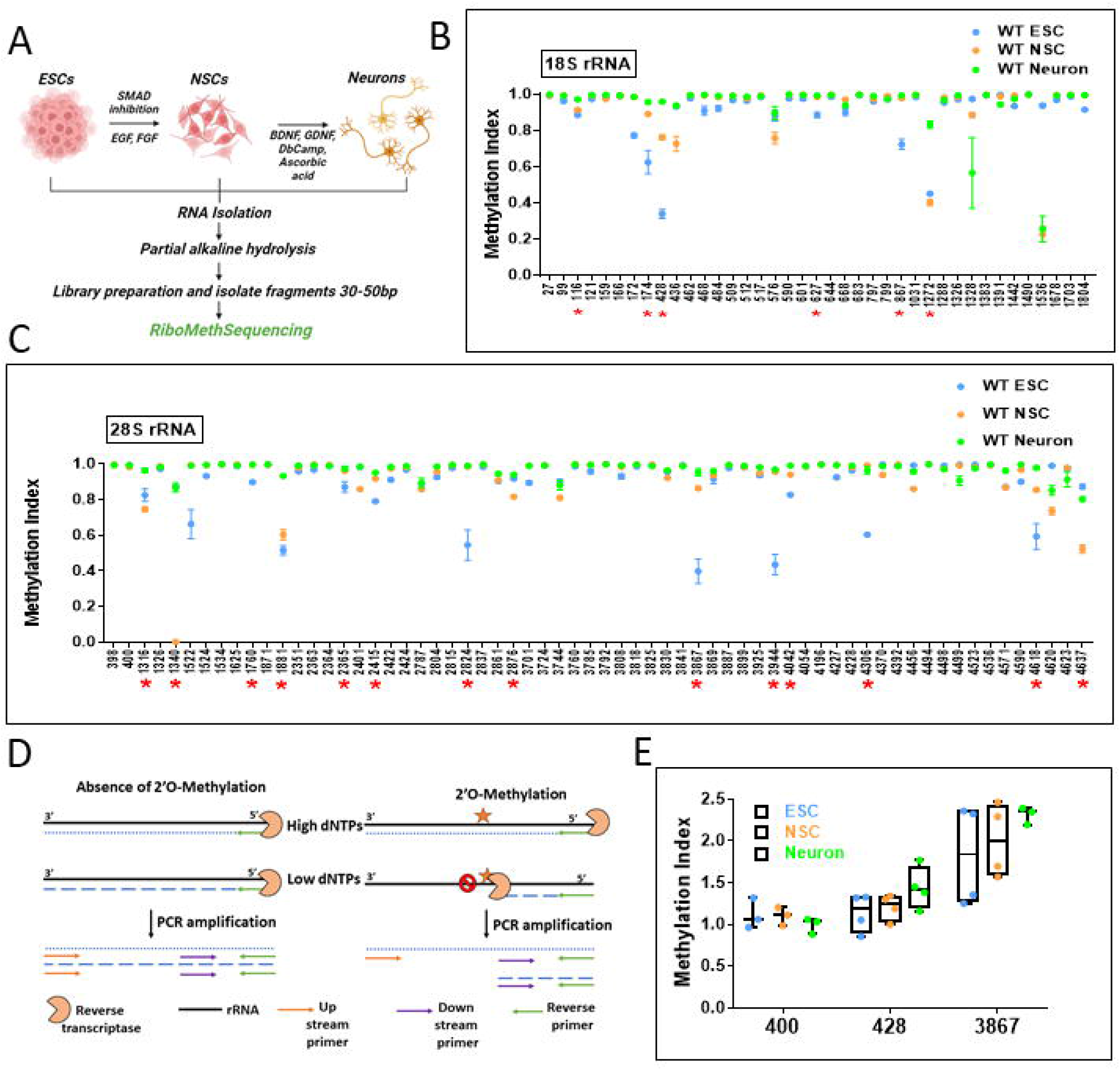

Next, we independently validated the changes in 2’O-methylation on specific sites of 18S and 28S rRNA through a qPCR-based tool referred to as RTL-P ^17^. Details of the primer design and product amplification have been described in **Figure 1D**. We have validated the changes in methylation for three different positions on rRNA-site 428 in 18S rRNA and sites 400 and 3867 in 28S rRNA. Position 391 on 28s rRNA was found to be methylated in all 3 differentiation stages (**Figure 1C**). Sites 428 and 3867 in 18S and 28S rRNA respectively show increasing trends of methylation in the NSC and neuronal stages (**Figure 1C**). Using RTL-P, we confirmed the complete methylation of Site 400 across the 3 different cell types and increased methylation of sites 428 and 3867 as the cells differentiated into neurons, validating the RMS data generated by next-generation sequencing (**Figure 1E**). Together, our results show maximum hypomethylation of rRNA in the ESCs which significantly decreases as the cells differentiate to post mitotic neurons.

### The effect of FMRP on 2’O-methylation of rRNA is maximum in ESCs

Our published work demonstrated a novel role for FMRP in regulating the methylation of 2’O ribose sugars of specific bases in ESCs^11^. In the current study, we aimed to understand the effect of FMRP on 2’O-methylation of rRNA during the differentiation of ESCs into NSCs and cortical neurons. For this, we used H9 ESC and FMR1 KO H9 ESC lines. The knockout of FMRP was performed through CRISPR-Cas9 deletion of exon1 of the FMR1 gene **(Figure S2D**)^18^. FMR1 KO ESCs were characterized for stem cell markers OCT4 and Nanog (**Figure 2A and S2A**). ESCs were differentiated into NSCs and neurons as described earlier^19^. Differentiated states were confirmed by the presence of Nestin and Pax6 in NSCs and MAP2 and VGlut1in neurons (**Figure S2B and 2C**).

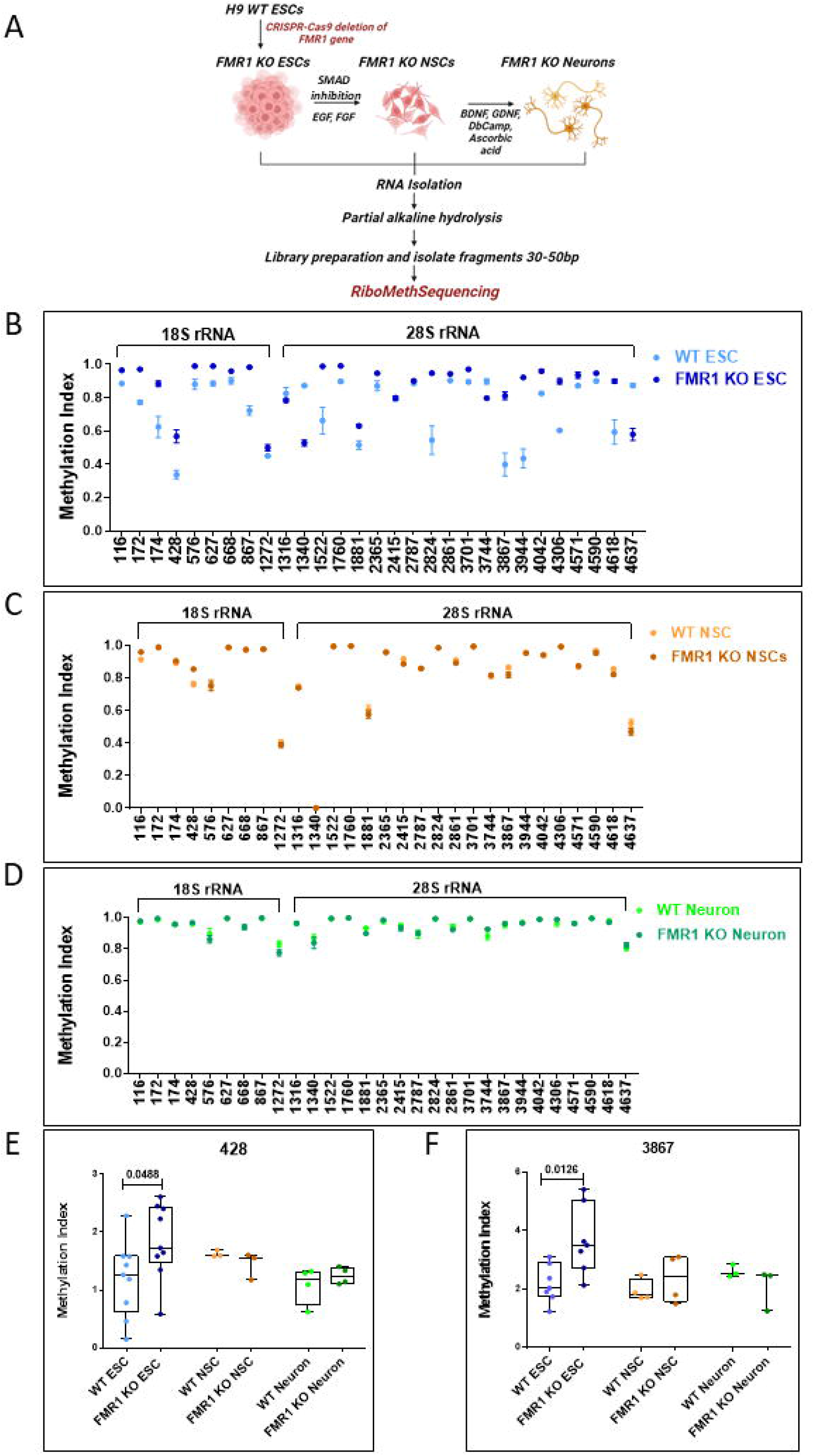

To understand how FMRP influences 2’O-methylation levels at each stage of differentiation, we performed RiboMethSequencing from ESCs, NSCs, and neurons from both WT and FMR1 KO cell lines (**Figure 2A)**. RMS data from FMR1 KO cells indicates maximum hypomethylation of rRNA was in the ESC stage compared to NSCs and neurons, which was similar to our observation in the WT condition. This suggests that the overall trend of increasing 2’O-methylation among ESC, NSC, and mature neurons does not change between WT and FMR1 KO conditions. (**Figure 2B, Figure S2E and S2F**). However, to study specific changes in 2’O-methylation status due to the loss of FMRP, we selected sites in WT ESC 18S and 28S rRNA that have an MI score less than/ equal to 0.9 and examined their MI in the FMR1 KO ESCs (**Figure 2B**). Furthermore, we compared the MI of these sites in WT and FMR1 KO NSCs (**Figure 2C**) and neurons (**Figure 2D**). We observed that the fold difference in 2’O-methylation between the WT and KO conditions of certain sites (e.g. Site 428 in 18S rRNA and site 2824 in 28S rRNA) drastically reduces from the ESC to NSC to neuronal types. Additionally, the number of hypomethylated sites (6 sites in 18S rRNA and 12 sites in 28S rRNA) reduced as we differentiated ESCs into NSCs and neurons. Details of these sites are provided in **Table 1** (Highlighted in red).

Further, we plotted a heat map by grouping variable positions and saturated positions across the differentiated cell states and compared them with the FMR1 KO condition (**Figure S2G and S2H**). We have indicated the sites on 18S and 28S rRNA showing significant changes in 2’O-methylation between WT and FMR1 KO ESCs/NSCs/neurons in **Tables 3-5**. These results indicate that the number of hypomethylated positions in both 18S and 28S rRNA are reduced over differentiation suggesting that the effect of FMRP on rRNA 2’O-methylation is maximum in ESCs. To further validate this result, we selected positions 428 from 18S rRNA and 3867 from 28S rRNA. The Methylation Index for these positions was measured by RTL-P in WT and FMR1 KO ESCs, NSCs, and neurons (**Figure 2E and Figure 2F**). In ESCs, sites 428 and 3867 show a significant increase in methylation status in the absence of FMRP compared to the WT condition. Further validation of the same sites in NSCs and neurons shows no difference in 2’O-methylation between WT and FMR1 KO conditions, suggesting that the role of FMRP in regulating 2’O-methylation is reduced across neuronal differentiation (**Figure 2E and Figure 2F**).

### rRNA hypermethylation in NSCs is a result of reduced FMRP-snoRNA interaction

From our results, we observe a trend of hypermethylation in 18S and 28S rRNA as ESCs differentiate from NSCs. We have previously shown that FMRP interacts with C/D Box snoRNAs and regulates the 2’O-methylation profile of rRNAs in ESCs^11^. Hence, we wanted to investigate the effect of FMRP-snoRNA interaction on 2’O-methylation in the context of neuronal differentiation^11^. To begin with, we examined the relative expression of selected C/D Box snoRNAs in ESCs and NSCs. We chose snoRNA candidates based on the sites that were hypomethylated on 18S and 28S rRNA in WT ESCs. We observed that the steady-state expression of these C/D Box snoRNAs did not significantly alter as ESCs differentiate into NSCs **(Figure 3A)**. Further, we observed no significant difference in the levels of the target snoRNAs between WT and FMR1 KO ESCs and NSCs confirming that FMRP does not affect the steady-state expression of these snoRNAs **(Figure 3B and Figure S3A)**.

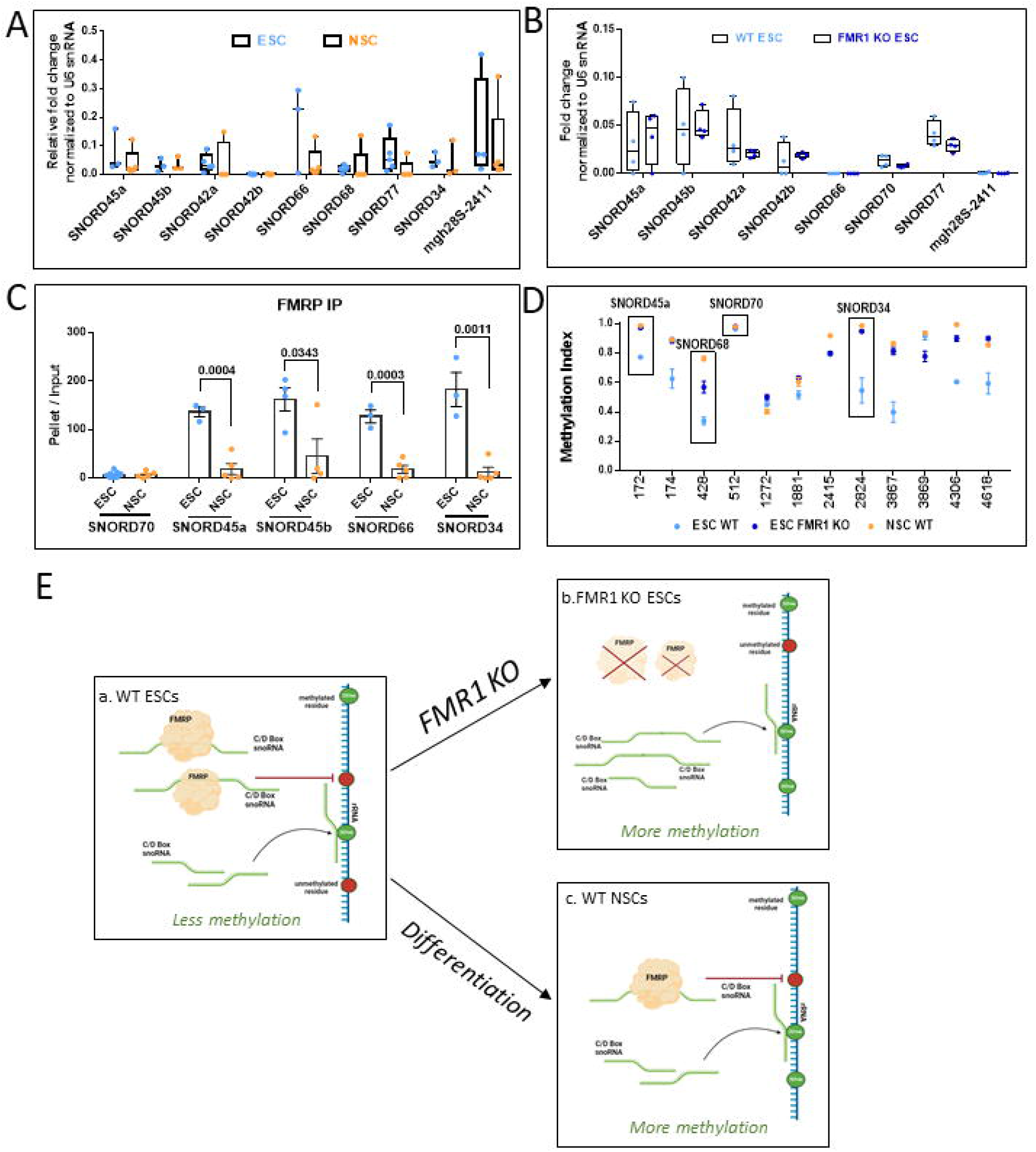

Since we did not capture any alterations in the levels of snoRNAs along differentiation or in the FMR1 KO condition, we investigated whether the altered 2’O-methylation pattern between WT ESCs and WT NSCs could be due to differences in FMRP-snoRNA interactions. Our previous work indicated that FMRP binds to several C/D box snoRNAs in ESCs as well as in NSCs^11^. However, the extent of FMRP-snoRNA interaction between these two cell states was not known. To test this, we performed an FMRP-immunoprecipitation from ESCs and NSCs and quantified the copy number of specific snoRNAs that were bound to FMRP in these two systems through qPCR **(Figure 3C)**. We observed that the extent of binding of selected snoRNAs to FMRP is significantly reduced in WT NSCs in comparison to WT ESCs **(Figure 3C)**.

We mapped the FMRP-bound snoRNAs to their respective target sites on rRNA to examine the changes in 2’O-methylation between WT ESCs, FMR1 KO ESCs, and WT NSCs **(Figure 3D)**. We observe that the sites corresponding to FMRP-bound snoRNAs shift from hypomethylation to hypermethylation state between WT and FMR1 KO ESCs **(Figure 3D)**. Similarly, we see a similar shift from hypomethylation to hypermethylation when we compare WT ESCs and WT NSCs for these sites **(Figure 3D)**. Thus, FMRP has a strong affinity to selected C/D Box snoRNAs in the WT ESCs which results in the hypomethylation of the sites targeted by these snoRNAs **(Figure 3E)**. This interaction is lost in FMR1 KO ESCs or is reduced in the case of WT NSCs, both of which result in hypermethylation of the sites targeted by the FMRP-bound snoRNAs **(Figure 3E and Table 4)**.

### Loss of FMRP results in protein synthesis defects in ESCs but not in NSCs

Cell state transitions are controlled by changes in global protein synthesis. To capture changes in global protein synthesis in the absence of FMRP along the differentiation of ESC to NSC, we made use of a non-canonical amino acid tagging system called FUNCAT^20^. The rate of production of newly synthesized proteins was measured in WT and FMR1 KO ESCs and NSCs through the quantification of the FUNCAT signal, which was normalized to endogenous α-tubulin protein levels in each condition (**Figure 4A**). We observed that isolated ESCs lose their pluripotency signal upon separation from the ESC colony. Hence we quantified the total FUNCAT and α-tubulin signal from whole ESC colonies and not from individual cells as we did in the case of NSCs. Our results indicate that the absence of FMRP caused a significant upregulation of global protein synthesis in ESCs compared to the WT condition (**Figure 4B and 4C**). Interestingly, we did not capture this trend in the differentiated NSCs (**Figure 4D and 4E**). There was no significant difference observed in the rates of translation between WT and FMR1 KO NSCs (**Figure 4D and 4E)**. This finding indicates that FMRP might have a prominent role in regulating translation at early developmental stages as opposed to intermediate stages of differentiation. To confirm our observations, we also measured rates of global protein synthesis in WT and FMR1 KO ESCs by quantifying the levels of puromycin incorporation between the two conditions (**Figure 4F**). We observe a significant increase in the levels of puromycin-labelled proteins in the absence of FMRP (FMR1 KO ESCs) indicating an overall increase in translation (**Figure S4A-S4D**). Since we hypothesize that increased 2’O-methylation on rRNA could result in increased rates of translation and WT NSCs have hypermethylated rRNA residues compared to WT ESC rRNA, we examined the rates of protein synthesis between WT ESCs and WT NSCs by measuring the levels of puromycin-labeled proteins. We did not measure this parameter through FUNCAT since it is not possible to compare the FUNCAT intensity between ESC colonies and individual NSCs. Our data shows that there is a significant increase in puromycin incorporation in WT NSCs compared to WT ESCs indicating that overall protein synthesis increases as cells differentiate from ESCs to NSCs (**Figure 4G-H and Figure S4E-F**).

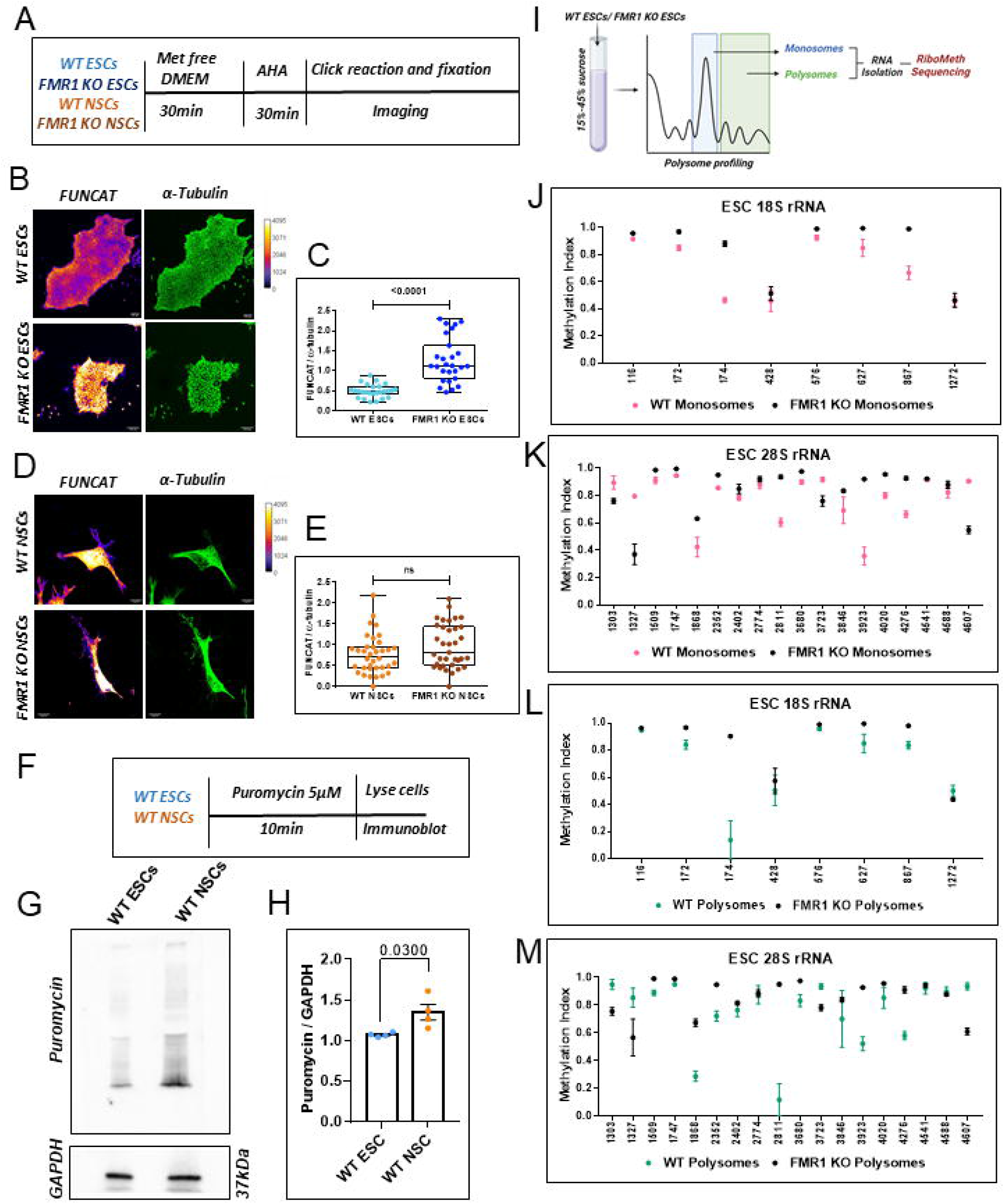

From our previous study, we know that FMRP regulates translation by affecting the epitranscriptome of the ribosome ^11^. We aimed to understand the effect of FMRP on the 2’O-methylation of translating monosomes and polysomes. For this, total cell lysate from WT and FMR1 KO ESCs was loaded on a linear sucrose density gradient and components were separated based on density through ultracentrifugation. We collected RNA from the pools of monosomes and Polysomes from WT and FMR1 KO conditions and subjected the RNA to RiboMethSequencing (**Figure 4I)**. Our data suggests that there are more hypomethylated sites in the 28S rRNA compared to 18S rRNA in both monosome and polysome populations **(Figure 4J-M and S4G-J)** and the absence of FMRP results in the hypermethylation of a majority of these sites in both the ribosomal populations (**Figure 4J-M and S4G-J)**.

## Discussion

Cellular differentiation is an event where a state of specialization is achieved to facilitate a unique cellular function. This process is contributed by an amalgamation of various processes such as transcription, epigenetic changes epitranscriptomic changes, protein synthesis, and cell signaling. Our study focuses on specialized ribosomes generated from epitranscriptome changes, which provide an important layer of complexity to protein synthesis during neuronal differentiation of pluripotent embryonic stem cells. Ribosomal RNA heterogeneity is generated primarily through altered sequence or epitranscriptomic modification of the rRNA ^7,8^. Our study shows a distinct pattern of 2’O-methylation on both 18S and 28S rRNA in human ESCs. We observe that many of these sites on rRNA are hypomethylated in the stem cell state. However, these same sites get further methylated when the cells are differentiated along the neuronal lineage. As cells achieve their post-mitotic fate, we observe the highest number of completely methylated sites. In other words, there is more 2’O hypomethylation of rRNAs in ESCs which reduces as ESCs are differentiated into NSCs and neurons (**Figure 5B)**. This is a very surprising result since neurons are highly polarized cells that will require elaborate compartmentalized and activity-mediated protein synthesis. Therefore, we expected a higher level of specialized ribosomes in them. On the contrary, our results indicate the highest level of rRNA 2’O hypomethylation is in ESCs and relatively reduced 2’O-methylation is in neurons. While considering our results, it is important to note that our RMS was performed with lysates from whole neurons and not from specific compartments. Neurons show localized protein translation at the synapses, tightly regulated by synaptic activity^21–23^. Hence, it is possible that there could exist a higher level of hypomethylated ribosomes within these compartments. Broadly, our data suggests that ESCs possess higher rRNA hypomethylation which we correlate with the pluripotent state of the cell.

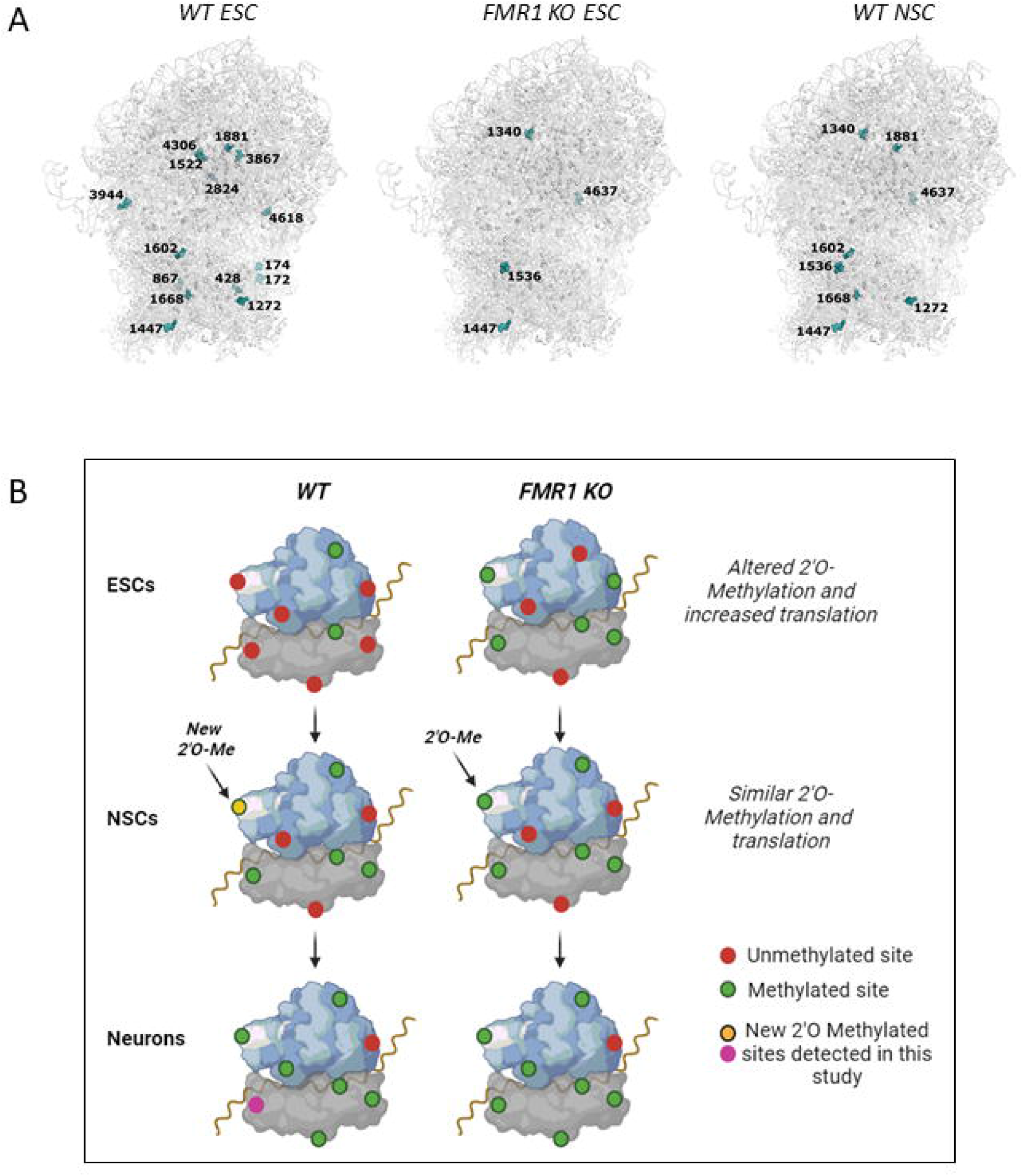

We observe a maximum number of hypomethylated sites in ESCs indicating a very high number of partially methylated ribosomes. This seemingly counterintuitive finding becomes logical once we carefully consider the pluripotent nature of ESCs. Translation rate is presumably low in ESCs and is thought to go up as they differentiate^1^. Cell differentiation is a highly dynamic process that occurs in response to various intrinsic and extrinsic cues. This requires quick proteomic remodeling which is largely determined by translation regulation. It is essential for ESCs to rapidly respond to these cues either by enhancing transcription or by priming the existing ribosomes. Here, we argue that a high level of hypomethylated ribosomes provides ESCs with such a potential. The high level of hypomethylated rRNA in ESCs suggests that there are large numbers of different ribosomal pools, each having its distinct pattern of methylation (**Figure 4J-M**). We argue that since ESCs are primed to differentiate into multiple germ layers, the system is equipped to translate different pools of mRNAs when required. In other words, we propose that pluripotency in ESCs is maintained because of the multiple pools of specialized ribosomes. Once differentiation is initiated, rRNA gets hypermethylated and the cells begin to produce a more homogenous population of ribosomes.

Previously we have shown that FMRP associates with a specific set of C/D box snoRNAs in ESCs and this can lead to the generation of specific rRNA methylation patterns and thus specialized ribosomes ^11^. Our current data shows that the steady-state expression of specific snoRNAs is unaltered between ESCs and NSCs. Further, the expression of specific snoRNAs is not affected in the absence of FMRP (**Figure 3B and Figure S3A**). This finding was anticipated as snoRNAs are very abundant yet essential for the process of ribosome biogenesis^24^. However, this result did not explain the differences in 2’O-methylation that we captured between ESCs and NSCs. Further, this also did not explain the trend of hypermethylation we observed in conditions where FMRP is absent. We hypothesize that the differential methylation of rRNA between cell states could be due to the differential interaction of FMRP with its target snoRNAs. We have shown that the interaction of FMRP with its target C/D Box snoRNAs is similar across different ESC lines such as Shef4 ESCs and H9 ESCs^11^. Our current data shows that the extent of FMRP-snoRNA interaction is significantly reduced from ESCs to NSCs (**Figure 3C**). This suggests that the binding of FMRP to snoRNAs is the strongest in ESCs making them less available to target the methylation sites on rRNA (**Figure 3D**). This data indicates that FMRP-dependent snoRNAs may be a critical factor in defining specialized ribosomes during neuronal differentiation. The exact molecular reasoning for this is unclear. Due to the higher sequestration of snoRNAs by FMRP in ESCs, we capture a high number of hypomethylated sites in ESC ribosomes. Since this interaction is reduced in NSCs, an increase in the number of available guide snoRNAs leads to more methylation on NSC rRNA **(Figure 3E and Figure 5B**). The decrease in FMRP-mediated snoRNA sequestration between WT ESCs and WT NSCs is very similar to the loss of snoRNA sequestration in FMR1 KO ESCs (**Figure 3E**). Correspondingly, the availability of free FMRP-target snoRNAs in FMR1 KO ESCs leads to the hypermethylation of rRNA (**Figure 3E and Figure 5B**). Hence, we can conclude that a shift in the extent of snoRNA sequestration by FMRP can determine the impact on rRNA methylation.

Since there is mounting evidence to link epitranscriptomic changes in rRNA to altered translation^25,26^, we decided to test the effect of altered rRNA 2’O-methylation on global protein synthesis in the ESCs and NSCs in the presence or absence of FMRP. FMRP is a regulator of translation, however, it is predominantly known to inhibit translation through the stalling of ribosomes^27,28^. Our analysis indicates that FMR1 KO ESCs show a significant upregulation of overall protein synthesis compared to WT H9 ESCs (**Figures 4B and 4C**). This phenomenon is only evident at the ESC stage. There was no difference in the global protein synthesis between WT-NSCs and FMR1 KO NSCs. Interestingly, the correlating observation of rRNA hypermethylation to increased protein synthesis was captured when we compared ESCs with their differentiated neuronal precursor forms (**Figure 4G and 4H**). Since protein expression during cellular differentiation is largely controlled by ribosomal function, this regulation is likely determined by cell-state-specific specialized ribosomes. Currently, we do not know the consequence of a 2’O Methylated rRNA residue on protein synthesis. However, it is clear that rRNA hypermethylation increases translation and hypomethylation reduces translation.

In addition, our general observation shows that 28S rRNA possesses the maximum number of hypomethylated residues in monosomal and polysomal pools in comparison to the 18S rRNA. This result is in line with the concept that the rRNA of the ribosomal small subunit is less variable compared to the large subunit^29,30^. Also, the absence of FMRP causes hypermethylation of a majority of sites across 18S and 28S rRNA in both the ribosomal populations (**Figure 5B)**. This implies that the occurrence of specialized ribosomes and their alteration due to the loss of FMRP is only significant at the early stages of embryonic development (**Figure 5B**). Further, the importance of FMRP in regulating ribosome biogenesis might be particularly relevant at the ESC state while FMRP might adopt alternate roles in the NSCs and neurons to regulate protein synthesis.

In summary, our study shows a clear role of FMRP-dependent ribosome heterogeneity in ESCs which is supported by our translation assays and snoRNA interaction. This will be the first report to show how RNA-binding proteins like FMRP may contribute to generating differential 2’O-methylation across the differentiation of pluripotent cells to terminal differentiated cells. Our data suggests that the key methylation positions vary between (WT and FMR1 KO ribosomes and along differentiation around the PTC (Peptidyl Transferase Center) (**Figure 5A**) and studying them in detail would open up many interesting avenues to understand 2’O-methylation-dependent translation regulation. Further, our study also shows how distinct 2’O-methylation patterns on rRNA can be used as indicators of specific cell states during differentiation and development.

## Materials and Methods

### Ethics statement

All the human stem cell work was carried out as per approval from the Institutional Human Ethics Committee and Institutional Biosafety Committee at InStem, Bengaluru, India, and Centre for Brain Research, Indian Institute of Science Campus, Bangalore, India.

### Embryonic Stem Cell culture

H9 ESCs and FMR1 KO ESCs were cultured on Matrigel (#3545277 BD Biosciences) coated plates containing mTeSR1 medium (#5850, StemCell Technologies) at 37°C in a 5% CO2 environment. Cells were passaged with an enzyme cocktail containing 1 mg/ml of Collagenase type IV (#17104019, Invitrogen), 20% KOSR (#10828010, Gibco), 0.25% Trypsin, and 1 mM CaCl2 dissolved in 1X PBS without CaCl2 or MgCl2 pH 7.2. For immunostaining experiments, H9 ESC colonies were plated on Matrigel-coated glass coverslips and cultured as mentioned above. H9 ESCs were further differentiated into Neural Precursor Cells (NPCs) by inducing them with a Neural Induction Medium for 14 days^19^. The protocol for neural differentiation was adapted from *Shi et al*^19^to differentiate iPSCs into forebrain glutamatergic neurons. The Neural Basic Media (NBM) for differentiation contained 50% DMEM F-12 (21331–020, ThermoFisher Scientific), 50% Neurobasal, 0.1% PenStrep, Glutamax, N2 (17502–048, ThermoFisher Scientific), and B27 without Vitamin A (12587–010, ThermoFisher Scientific). Once the iPSCs reached 70-80% confluency, they were subjected to monolayer neural induction by changing the mTeSR1 media to Neural Induction Media (NIM). NIM is composed of NBM supplemented with small molecules SB431542 (10 µM, an inhibitor of TGFβ pathway) (72232, Stem Cell Technologies) and LDN193189 (0.1 µM, an inhibitor of BMP pathway) (72142, Stem Cell Technologies). The cells were subjected to neural induction for 12-15 days by changing the media every day till a uniform neuroepithelial had formed. After the induction, the monolayer was dissociated using Accutase (A6964, Sigma), and centrifuged at 1200 rpm for 3 minutes at room temperature. The cells were plated overnight in NIM containing 10 µM ROCK inhibitor (Y0503, Sigma) on pre-coated poly-L-ornithine/laminin dishes. Poly-L-Ornithine (1:10 dilution in 1X PBS) (P4957, Sigma) coating was performed at 37°C for a minimum of 4 hours, washed thrice with 1X PBS, followed by overnight coating with Laminin (5 µg/ml diluted in 1X PBS) (L2020, Sigma) at 37°C. The NSCs were maintained in Neural Expansion Media (NEM) composed of NBM supplemented with FGF (10 ng/ml) (100-18C, Peprotech) and EGF (10 ng/ml) (AF-100-15, Peprotech). Neuronal maturation and terminal differentiation were achieved by plating the NSCs at a density of 25,000-35,000 cells/ cm^2^ in the Neural Maturation Media (NMM) composed of NBM supplemented with BDNF (20 ng/ml) (450-02, Peprotech), GDNF (10 ng/ml) (450-10, Peprotech), L-Ascorbic Acid (200 µM) (A4403, Sigma) and db-Camp (50 µM) (D0627, Sigma). The neurons were subjected to maturation for 4-5 weeks by supplementing them with NMM every 4-5 days.

### Characterization of stem cells

ESCs, NSCs, and neurons were fixed with 4% PFA for 15 minutes followed by 1X PBS wash and permeabilization with 0.3% Triton X-100 (made in TBS_50_) for 10 minutes. This was followed by 1 hour blocking with 2% BSA and 2% FBS prepared in TBS_50_T (with 0.1% Triton X-100). They were incubated with the primary antibody (prepared in blocking buffer) overnight at 4°C. This was followed by 3 washes with TSB_50_T and 1-hour incubation with the secondary antibody (prepared in blocking buffer) at room temperature. After 3 washes with TBS_50_T, the cells were mounted with Mowiol.

### Metabolic labeling

ESCs and NSCs were incubated in methionine-free Dulbecco’s Modified Essential Medium (Thermo# 21013024) for 30min followed by the addition of azidohomoalanine (AHA; 1μM #C10102, Thermo) in the same medium. This was incubated for 30 minutes and fixed with 4%PFA for 10 minutes. Cells were then permeabilized in PBS+0.3% Triton X-100 solution and blocked with buffer containing PBS+0.1% Triton X-100 + 2%BSA + 4% FBS solution. Newly synthesized proteins were then labeled with Alexa-Fluor-555–alkyne [Alexa Fluor 555 5-carboxamido-(propargyl), bis (triethylammonium salt) (#A20013, ThermoFisher scientific], by allowing the fluorophore alkyne to react with AHA azide group through click chemistry(CLICK-iT cell reaction buffer kit, #C10269). The cells were subjected to immunostaining for a-tubulin (ESCs and NSCs) and MAP2 (neurons) to identify the cells. Mowiol® 4-88 mounting media was used to mount the coverslips (#81381 Sigma).

### Imaging

Mounted coverslips were imaged on an Olympus FV3000 confocal laser scanning inverted microscope with a 20X objective. The pinhole was kept at 1 Airy Unit and the optical zoom at 2X to satisfy Nyquist’s sampling criteria in XY direction. The objective was moved in Z-direction with a step size of 1 μM (∼8-10 Z-slices) to collect light from the planes above and below the focal plane. For FUNCAT, the cells were identified using α-tubulin channel respectively. The image analysis was performed using ImageJ software and the maximum intensity projection of the slices was used for quantification of the mean fluorescent intensities. The region of interest (ROI) was drawn around the cells using the a-tubulin channel. Data is represented as box plots indicating the quantification of the FUNCAT fluorescent intensity normalized to α-tubulin fluorescent intensity. The box extends from the 25th to the 75th percentile. The middlemost line represents the median of the dataset and the whiskers of the box plot range from minimum to maximum data points.

### Linear sucrose density centrifugation

Polysome assay was done from WT and FMR1 KO ESC lysate as described previously^31^. In brief, cell lysate was separated on a 15%–45% linear sucrose gradient in the presence of 0.1mg/ml Cycloheximide (CHX) (#C7698-5G, Sigma) and Phosphatase inhibitor (#4906837001, Roche) by centrifugation at 39,000 rpm in SW41 rotor for 90 min. The sample was fractionated into 12 1.0 mL fractions with continuous UV absorbance measurement (A254). Fractions were pooled as monosomes (F4 and F5) and polysomes (F6-12) according to ribosomal subunit distribution based on the peaks. RNA was isolated from the pooled fractions and subjected to RiboMethSequencing and RTL-P.

### snoRNA quantification

CDNA of snoRNA was prepared using reverse primers specific to individual snoRNA candidates ^11^. CDNA was amplified using SYBR premix by qPCR. Arbitrary copy numbers were calculated from a standard curve drawn from Ct values obtained from serial dilutions of cDNA for snoRNA candidate HBII99. Copy numbers for various snoRNA candidates were obtained using the equation generated from the standard curve.

### Immunoprecipitation

ESCs and NSCs were lysed in 1% NP40 containing lysis buffer (20 mM Tris-HCl pH 7.5, 150 mM NaCl, 5mM MgCl2 with protease and RNase inhibitors) and spun at 18000 rcf (12500 rpm) for 20 minutes at 4°C. Precleared supernatant was used for immunoprecipitation with Protein G Dynabeads. 5μg of anti-FMRP antibody was coupled to the Protein G Dynabeads. Lysates were incubated with antibody-conjugated beads for 1h at RT following which RNA was isolated using Trizol.

### Immunoblotting

The lysates from WT and FMR1 KO ESCs, NSCs, and neurons were characterized by western blot for the expression of FMRP. Briefly, the denatured lysates were run on 10% resolving and 5% stacking acrylamide gels and subjected to overnight transfer onto the PVDF membrane. The blots were subjected to blocking for 1h at room temperature using 5% Blotto prepared in TBST (TBS with 0.1% Tween-20). This was followed by primary antibody (prepared in blocking buffer) incubation at RT for 3 hours. HRP-tagged secondary antibodies were used for primary antibody detection. The secondary antibodies (prepared in blocking buffer) were incubated with the blots for 1h at room temperature. Three washes of TBST solution were given after primary and secondary antibody incubation. The blots were subjected to chemiluminescent-based detection of the HRP-tagged proteins.

### RiboMethSequencing and analysis

2μg of total RNA extracted from WT and FMR1 KO ESCs, NSCs, neurons, and ribosomal fractions was used for library preparation. RNA was hydrolyzed with alkaline Tris buffer (pH 10) at 95°C for 5 minutes and ethanol precipitated. Isolated RNA was run on a 12% TBE PAGE gel and a band corresponding to 30-50 bp was excised out. Sequencing libraries were prepared using the TruSeq small RNA library preparation Kit from Illumina and were sequenced on the Hiseq2500 platform. FastQC (v0.11.5) was used to assess the quality of the 50bp reads across all the samples. Adapter sequences (TGGAATTCTCGGGTGCCAAGG) were trimmed using Cutadapt (v2.10). The trimmed reads were aligned to the reference rRNA sequences (ENST00000606783–18S rRNA & ENST00000607521–28S rRNA) using bowtie (v1.1.2) with default parameters in the end-to-end mode. The alignment files were sorted and indexed using samtools (v1.7), which were then used for counting the number of 5’ and 3’ read-ends that were mapped to each position on the reference rRNA using bedtools (v2.25.0). The 5’ counts were shifted up by one position and combined with the 3’ counts to ascertain the methylated positions in the reference sequence. Further, RiboMeth-Seq scores^32^ were calculated for all the known methylated positions (64 from 28S rRNA and 42 from 18S rRNA) using custom bash and awk scripts. Heatmaps of the score C from the various samples were plotted in R using the package ‘pheatmap’.

### RTL-P (Reverse transcription at low-dNTP concentration followed by PCR)

2 ng of sample RNA was used for cDNA preparation using reverse primers (10μM) specific to methylation sites under high dNTP (10 mM) and low dNTP (1 nM) concentrations. For real-time PCR, we adopted a method from (Dong et al., 2012)^17^. We have used two forward primers for a methylation site; one up-stream (P1) and one down-stream (P2) from the methylation site, along with a common reverse primer (P3). Amplification with these sets of primers would yield one product over the methylation site which will be the longer product and another will be within the methylation site and would yield a small-length product. The extent of methylation for a given site is calculated as a methylation score as previously described in *D’Souza et al 2019*^*11*^.

### Puromycin labeling

ESCs and NSCs were incubated with 5 µM Puromycin (Cat no: P8833-25MG, Sigma) for 10 minutes. and were lysed in buffer (20 mm Tris-HCl, 100 mm KCl, 5 mm MgCl_2_, 1% NP40, 1 mm DTT, 1× protease inhibitor cocktail, and 1× phosphatase inhibitor). The protein levels were quantified from precleared lysates using the BCA method (Cat no: 23227, ThermoFisher Scientific). 50µg of total protein was loaded for all samples on a 10% SDS polyacrylamide gel. Immunoblots were stained with ponceau to ensure the successful transfer of proteins. The blots were blocked in 5% BSA made in 1× TBST. The blots were incubated in puromycin antibody (Cat no: MABE343, Sigma; 1:10000) for 3 hours at RT followed by anti-mouse HRP antibody (Cat. No: A9044, Sigma; 1:10000) for 1 hour at room temperature. The same immunoblots were stripped of the puromycin antibody (62.5 mM Tris Buffer (pH 6.8), 2% SDS, 0.7% Beta Mercaptoethanol). This was followed by incubation with GAPDH (Cat no. 2118S, Cell Signaling Technologies; 1:5000) for 1 hour at room temperature and anti-rabbit HRP antibody (Cat. No: A0545, Sigma; 1:10000) for 1 hour at room temperature. The puromycin signal was normalized to the GAPDH levels.

### Statistical analysis

All statistical analyses were performed using Graph Pad Prism software. The normality of the data was tested using the Kolmogorov-Smirnov test. For experiments with less than 5 data points, parametric statistical tests were applied. Data were represented as mean ± SEM in all in-vitro and polysome experiment graphs. FUNCAT data was represented as boxes and whiskers with all the individual data points. Statistical significance was calculated using Unpaired Student’s t-test (2 tailed with equal variance) in cases where 2 groups were being compared. One-way ANOVA was used for multiple group comparisons, followed by Tukey’s multiple comparison tests, Bonferroni’s multiple comparison test, or Dunnett’s multiple comparison test. Unpaired Student’s t-test was used to calculate statistical significance for all snoRNA qPCR assays and Puromycin incorporation assays.

## Acknowledgments

The work has been funded by the Core funds from CBR, IISc. R.S.M. acknowledges the SERB-DST grant (Reference number: EMR/2016/006313). D.P. would like to acknowledge his DST Swarnajayanti Fellowship (DST/SJF/LSA-02/2015-16). N.K.C.G acknowledges his National Postdoctoral Fellowship from DST for the same (Reference number: PDF/2018/000663). N.H. would like to acknowledge CSIR and DBT-InStem for funding. We would like to thank the Stem Cell Facilities at InStem and CBR. We would also like to thank the Sequencing facility at InStem and the Imaging facility at CBR. We would like to thank all Muddashetty laboratory members for their suggestions and advice during this work.

## Author Contributions

Conceptualization - M.N.D., N.K.C.G., N.H, D.P and R.S.M.; Validation - M.N.D. and N.K.C.G.; Biochemical assays-SWQ; Formal Analysis – N.H, M.N.D and N.K.C.G.; Resources – D.P and R.S.M; Writing, Review & Editing, M.N.D., N.K.C.G., N.H, D.P, and R.S.M.; Funding-D.P and R.S.M.; Supervision, R.S.M and D.P.

## Data Availability

Sequencing data have been deposited at NCBI Genbank under the following identifiers.

BioProject Accession: PRJNA1129659

Reviewer Link: https://dataview.ncbi.nlm.nih.gov/object/PRJNA1129659?reviewer=oe0h4lu4haoqgg16132a2dt6f2

---

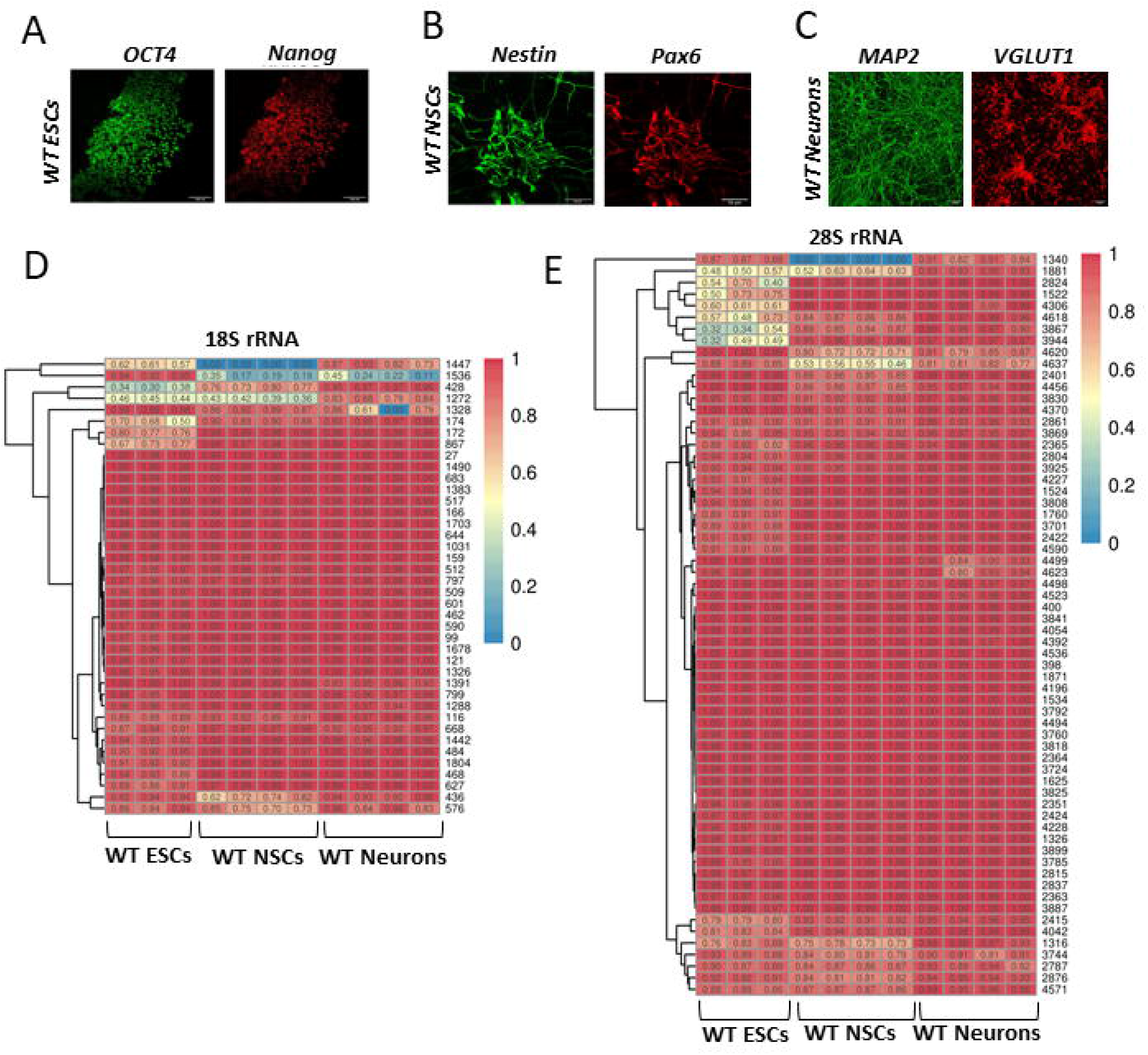

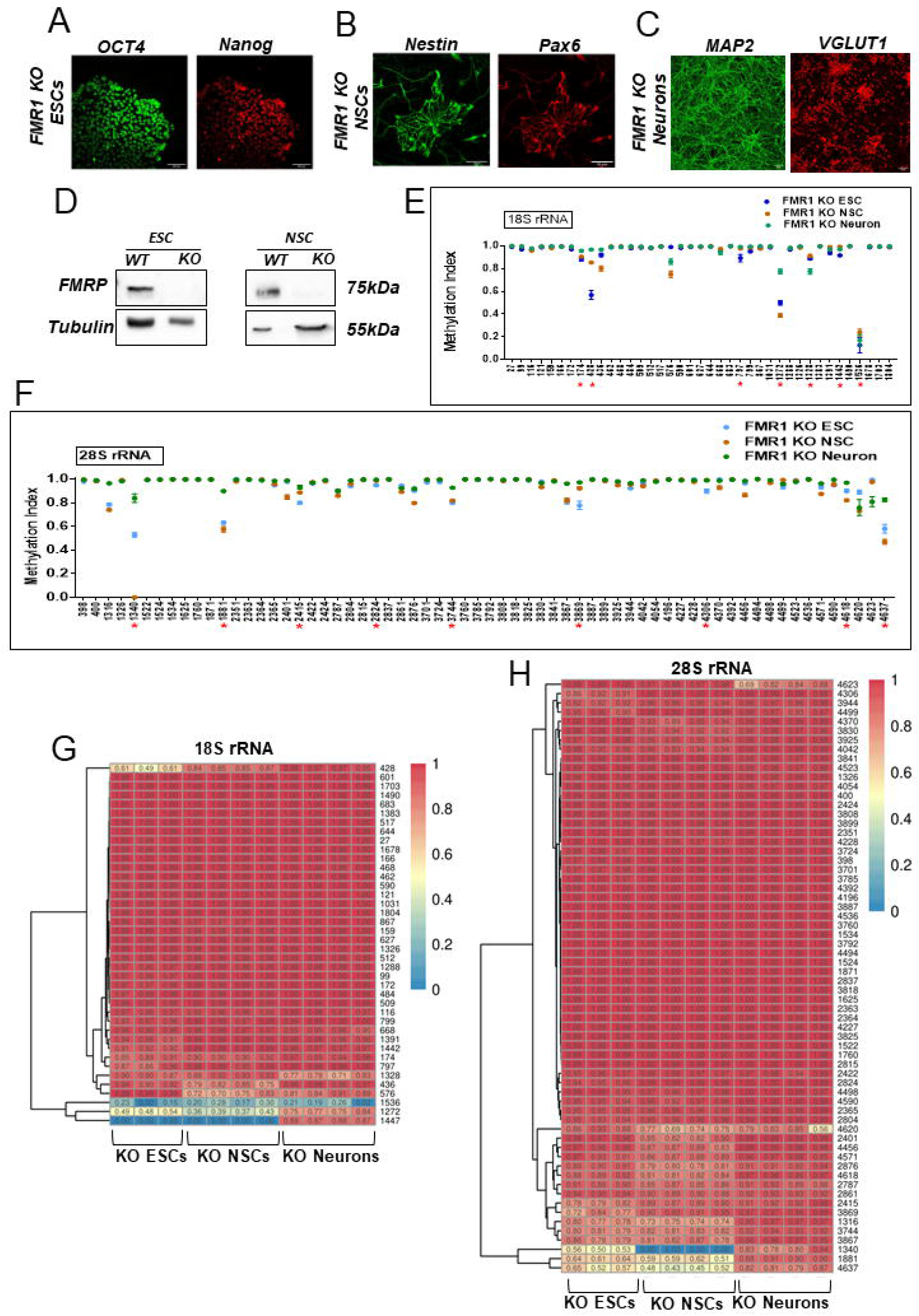

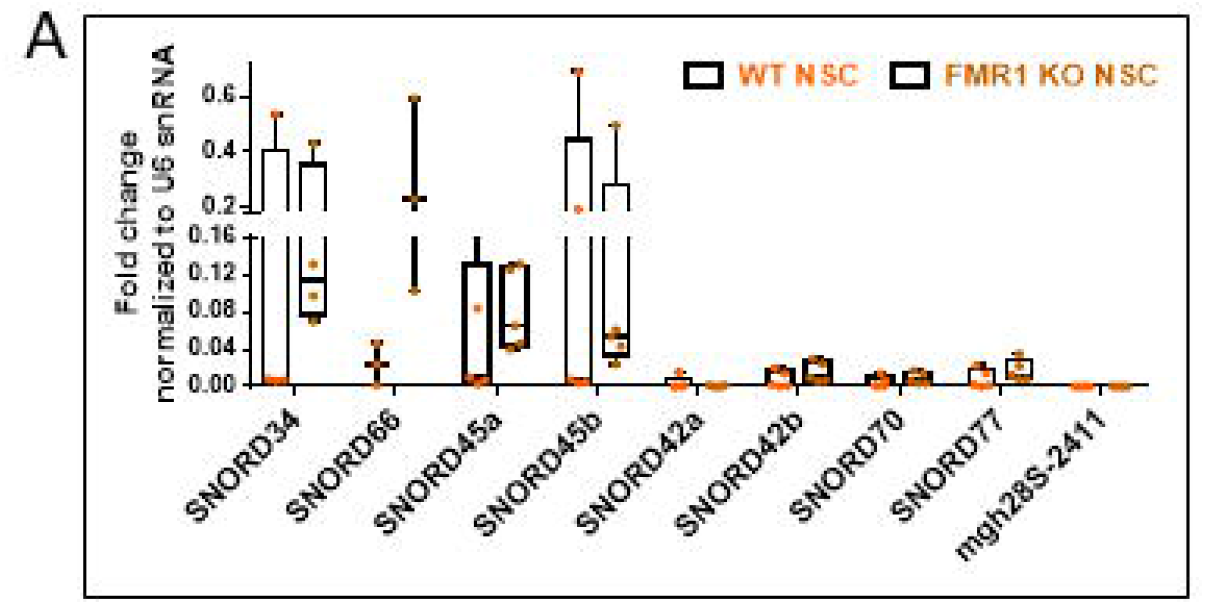

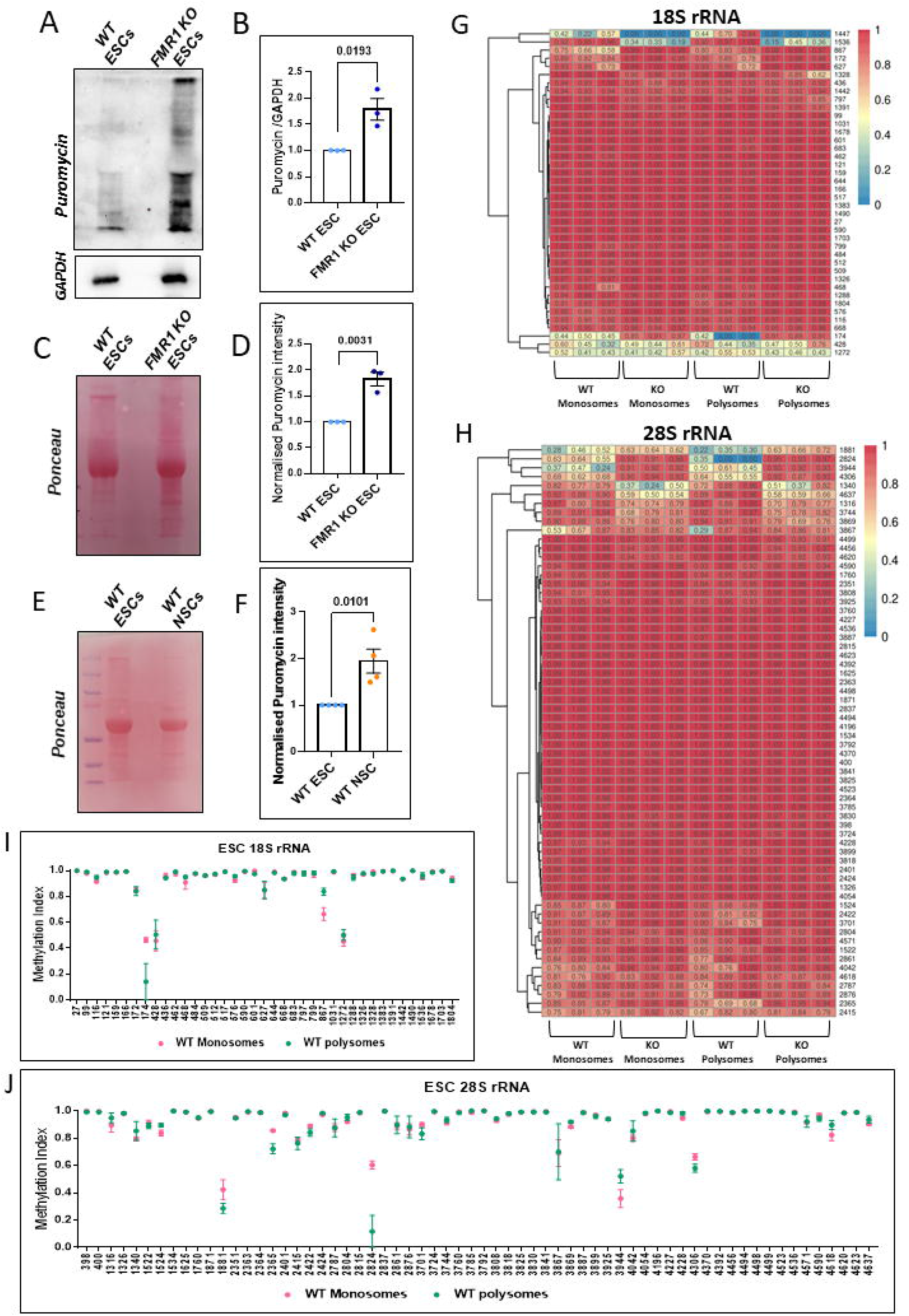

