## Supplementary material for "Altering rRNA 2’O-methylation pattern during neuronal differentiation is regulated by FMRP": Figure legends and Supplemental Tables

**Figure 1: Changes in 2’O-Methylation Pattern of rRNA along neuronal differentiation of H9 ESCs.**

A. Schematic for RiboMeth Sequencing from H9 ESCs, H9 NSCs, and H9 neurons.
B. Methylation Index of the sites on 18S rRNA in H9 ESCs, H9 NSCs, and H9 neurons. The x-axis represents the methylation position on 18S rRNA, and the y-axis represents the Methylation Index. Data is represented as Mean +/- SEM, n = 3 for ESCs, n=4 for NSCs and n=4 for neurons. *indicates sites that show an increase in 2’O-Methylation as ESCs are differentiated to NSCs and neurons.
C. Methylation Index of the sites on 28S rRNA in H9 ESCs, H9 NSCs and H9 neurons. The x-axis represents the methylation position on 28S rRNA, and the y-axis represents the Methylation Index. Data is represented as Mean +/- SEM, n = 3 for ESCs, n=4 for NSCs and n=4 for neurons. *indicates sites that show an increase in 2’O-Methylation as ESCs are differentiated to NSCs and neurons
D. Schematic of PCR-based RTL-P assay used to identify 2’O-methylation status of individual sites.
E. Box plot representing the changes in the Methylation Index for sites 400, 428, and 3867 between H9 ESCs, H9 NSCs, and H9 neurons measured through RTL-P. The box extends from the 25th to 75th percentile with the middlemost line representing the median of the dataset. Whiskers range from minimum to maximum data points. n=3-4 independent experiments. One-way ANOVA was performed for a given site and the multiple comparison test was performed with its values in ESC, NSC, and neuron state.

**Figure 2: Altered 2’O-Methylation pattern along neuronal differentiation in the absence of FMRP.**

A. Schematic showing the generation of FMR1 KO H9 ESCs through CRISPR-Cas9. rRNA from FMR1 KO ESCs, FMR1 KO NSCs, and FMR1 KO neurons were further processed for RiboMethSequencing.

B. Methylation indices of selected sites (MI> 0.90) on 18S rRNA and 28S rRNA in H9 WT and FMR1 KO ESCs. The x-axis represents the methylation position on 28S rRNA, and the y-axis represents the Methylation Index. Data represented as Mean +/- SEM, n = 3

C. Methylation indices of selected sites (**from B)** on 18S rRNA and 28S rRNA in H9 WT and FMR1 KO NSCs. The x-axis represents the methylation positions on 18S and 28S rRNA, and the y-axis represents the Methylation Index. Data represented as Mean +/- SEM, n = 4

D. Methylation indices of selected sites (**from B**) on 18S rRNA and 28S rRNA in H9 WT and FMR1 KO neurons. The x-axis represents the methylation positions on 18S and 28S rRNA, and the y-axis represents the Methylation Index. Data represented as Mean +/- SEM, n = 4

E. Box plot representing the change in Methylation Index for site 428 between WT and FMR1 KO ESCs, NSCs, and neurons measured through RTL-P. The box extends from the 25th to 75th percentile with the middlemost line representing the median of the dataset. Whiskers range from minimum to maximum data points. Unpaired t-test, n=9 for ESCs, n=3 for NSCs and n=4 for neurons

F. Box plot representing the change in Methylation Index for site 3867 between WT and FMR1 KO ESCs, NSCs, and neurons measured through RTL-P. The box extends from the 25th to the 75th percentile with the middlemost line representing the median of the dataset. Whiskers range from minimum to maximum data points. Unpaired t-test, n=9 for ESCs, n=3 for NSCs and n=4 for neurons.

**Figure 3: The interaction of FMRP with C/D Box snoRNA determines the extent of methylation**

A. Quantification of selected C/D box snoRNA expression in WT ESC and WT NSC. The graph represents fold change for a given snoRNA normalized to U6 snRNA. The box extends from the 25th to the 75th percentile with the middlemost line representing the median of the dataset. Whiskers range from minimum to maximum data points. Unpaired t-test, n=3-5

B. Quantification of C/D box snoRNA basal level expression in WT ESC and FMR1 KO ESC. The graph represents fold change for a given snoRNA normalized to U6 snRNA. The box extends from the 25th to the 75th percentile with the middlemost line representing the median of the dataset. Whiskers range from minimum to maximum data points. Unpaired t-test, n=4

C. Quantification of selected C/D box snoRNA enrichment in FMRP immunoprecipitates from WT ESC and WT NSC. The bar graph represents the pellet/input ratio for a given snoRNA normalized to U6 snRNA. Data represented as Mean +/-SEM. Unpaired t-test, n=3-9

D. Methylation indices of the sites on 18S and 28S rRNA in WT H9 ESCs which get hypermethylated due to a reduction in FMRP-snoRNA interaction (WT NSC) or loss of FMRP (FMR1 KO ESC). Boxed sites are targeted by FMRP-target snoRNA. The x-axis represents the methylation positions on 18S and 28S rRNA, and the y-axis represents the Methylation Index. n = 3. Data represented as Mean +/- SEM.

E. Model indicating the link between FMRP-snoRNA interaction and methylation pattern a) WT ESCs- FMRP sequesters C/D Box snoRNAs leading to rRNA hypomethylation. b) FMR1 KO ESCs- the loss of FMRP increases the availability of free C/D Box snoRNAs leading to rRNA hypermethylation and c) WT NSCs - reduced binding of FMRP to C/D Box snoRNAs leads to rRNA hypermethylation on selected sites resulting in a similar change in Methylation Index as FMR1 KO ESC condition.

**Figure 4: Altered 2'O-methylation affects translation in ESCs and NSCs.**

A. Schematic describing FUNCAT assay for the measurement of de-novo protein synthesis in WT and FMR1 KO ESCs and NSCs.

B. Representative images for FUNCAT and α-Tubulin fluorescent intensities in WT and FMR1 KO ESCs. (Scale bar—100 µm).

C. Box plot representing the quantification of the FUNCAT intensity normalized to α-Tubulin intensity for each colony of WT ESC and FMR1 KO ESC. The box extends from the 25th to the 75th percentile with the middlemost line representing the median of the dataset. Whiskers range from minimum to maximum data points. Unpaired t-test. n=30-35 colonies from 3 independent experiments.

D. Representative images for FUNCAT and α-Tubulin intensities in WT and FMR1 KO NSCs. (Scale bar—50 μm).

E. Box plot representing the quantification of the FUNCAT intensity normalized to α-Tubulin intensity for each cell of WT and FMR1 KO NSC. The box extends from the 25th to the 75th percentile with the middlemost line representing the median of the dataset. Whiskers range from minimum to maximum data points. Unpaired t-test. n=35-36 cells from 3 independent experiments.

F. Schematic describing Puromycin labeling of newly synthesized proteins to measure rates of translation in WT ESCs and WT NSCs.

G. Immunoblot indicating an increase in Puromycin incorporation (5μM for 10min at 37ºC) in WT NSCs compared to WT ESCs. 50μg of total protein was loaded per sample.

H. Quantification of Puromycin incorporation in WT ESCs and WT NSCs. Data is normalized to respective GAPDH levels. Mean +/- SEM from n=4 independent experiments.

I. Schematic for RiboMethSequencing from WT ESCs input, monosome, and polysome fractions after polysome profiling. WT ESCs were subjected to sucrose density gradient centrifugation on a 15-45% sucrose gradient. RNA was extracted from WT ESCs input, monosome, and polysome fractions and processed for RiboMeth Sequencing.

J. Methylation indices of the sites on 18S rRNA in WT ESC monosomes and polysomes. The x-axis represents the methylation positions on 18S rRNA, and the y-axis represents the Methylation Index. Data is represented as Mean +/- SEM, n = 3

K. Methylation indices of the sites on 28S rRNA in WT ESC monosomes and polysomes. The x-axis represents the methylation positions on 28S rRNA, and the y-axis represents the Methylation Index. Data represented as Mean +/- SEM, n = 3

L. Methylation indices of selected sites (MI>0.90) on 18S rRNA in ESC WT and FMR1 KO polysomes. The x-axis represents the methylation position on 18S rRNA, and the y-axis represents the Methylation Index, Data is represented as Mean +/- SEM, n = 3

M. Methylation indices of selected sites (MI>0.90) on 28S rRNA in ESC WT and FMR1 KO polysomes. The x-axis represents the respective methylation position on 28S rRNA, and the y-axis represents the Methylation Index, Data is represented as Mean +/- SEM, n = 3.

**Figure 5: Altering 2'O-Methylations on ribosomes during neuronal differentiation**

A. Front view of 2'O-methylation sites on WT ESC, FMR1 KO ESC, and WT NSC ribosomes. Hypomethylated sites are indicated in blue (MI <0.75).
B. Graphical abstract indicating altering patterns of 2'O-Methylation on rRNA across neuronal differentiation and in the absence of FMRP

**Supplementary Figure 1:**

A. Characterization of H9 ESCs with pluripotency marker OCT4 and Nanog (Scale bar—100 μm).

B. Characterization of H9 NSCs with differentiation marker Nestin and Pax6 (Scale bar—50 µm).

C. Characterization of H9 neurons with differentiation marker MAP2 and VGlut1 (Scale bar—50 µm).

D. Heat map indicating the Methylation indices of the sites on 18S rRNA in H9 ESCs, NSCs, and neurons. Heatmap indicates the hierarchical clustering of sites that show similar trends of changing methylation indices

E. Heat map indicating the Methylation indices of the sites on 28S rRNA in H9 ESCs, NSCs, and neurons.

**Supplementary Figure 2:**

A. Characterization of FMR1 KO ESCs with pluripotency marker OCT4 and Nanog (Scale bar—100 µm).

B. Characterization of FMR1 KO NSCs with differentiation marker Nestin and Pax6 (Scale bar—50 µm).

C. Characterization of FMR1 KO neurons with differentiation marker MAP2 and VGlut1 (Scale bar—50 µm).

D. Immunoblot indicating absence of FMRP in FMR1 KO ESCs and NSCs.

E. Methylation indices of the sites on 18S rRNA in FMR1 KO ESCs, FMR1 KO NSCs, and FMR1 KO neurons. The x-axis represents the methylation positions on 18S rRNA, and the y-axis represents the Methylation Index. Data represented as Mean +/- SEM, n = 3 for ESCs, n=4 for NSCs and n=4 for neurons. *indicates sites that show an increase in 2'O-Methylation as ESCs are differentiated to NSCs and neurons.

F. Methylation indices of the sites on 28S rRNA in FMR1 KO ESCs, FMR1 KO NSCs, and FMR1 KO neurons. The x-axis represents the respective methylation positions on 28S rRNA, and the y-axis represents the Methylation Index. Data represented as Mean +/- SEM, n = 3 for ESCs, n=4 for NSCs and n=4 for neurons. *indicates sites that show an increase in 2'O-Methylation as ESCs are differentiated to NSCs and neurons.

G. Heat map indicating the Methylation indices of the sites on 18S rRNA in FMR1 KO ESCs, FMR1 KO NSCs, and FMR1 KO neurons.

H. Heat map indicating the Methylation indices of the sites on 28S rRNA in FMR1 KO ESCs, FMR1 KO NSCs, and FMR1 KO neurons.

**Supplementary Figure 3:**

A. Quantification of C/D box snoRNA expression levels in WT NSCs and FMR1 KO NSCs. Box plots represent fold change for a given snoRNA normalized to U6 snRNA. The box extends from the 25th to the 75th percentile with the middlemost line representing the median of the dataset. Whiskers range from minimum to maximum data points. Unpaired t-test, n=5

**Supplementary Figure 4:**

A. Immunoblot indicating an increase in Puromycin incorporation (5µM for 10min at 37ºC) in WT ESCs and FMR1 KO ESCs. 50µg of total protein was loaded per sample.

B. Quantification of Puromycin incorporation in WT ESCs and FMR1 KO ESCs. Data is normalized to respective GAPDH levels. Mean +/- SEM from n=4 independent experiments.

C. Blot of WT ESC and FMR1 KO ESC lysate after puromycin incorporation stained with Ponceau.

D. Quantification of Puromycin incorporation in WT ESCs and FMR1 KO ESCs. Data is normalized to WT ESCs. Mean +/- SEM from n=4 independent experiments.

E. Blot of WT ESC and WT NSC lysate after puromycin incorporation stained with Ponceau.

F. Quantification of Puromycin incorporation in WT ESCs and WT NSCs. Data is normalized to WT ESCs. Mean +/- SEM from n=4 independent experiments.

G. Heat map indicating the Methylation indices of the sites on 18S rRNA in the monosome and polysome fractions of WT and FMR1 KO ESCs. N=3.

H. Heat map indicating the Methylation indices of the sites on 28S rRNA in the monosome and polysome fractions of WT and FMR1 KO ESCs. N=3.

I. Methylation Index of selected sites (MI>0.90) on 18S rRNA in ESC WT and FMR1 KO monosomes. The x-axis represents the respective methylation position on 18S rRNA, and the y-axis represents the fraction methylated, Data is represented as Mean +/- SEM, n = 3

J. Methylation Index of selected sites (MI>0.90) on 28S rRNA in ESC WT and FMR1 KO monosomes. The x-axis represents the respective methylation position on 28S rRNA, and the y-axis represents the fraction methylated, Data is represented as Mean +/- SEM, n = 3.

**Table 1:** Sites indicating significant shift from Hypomethylation to hypermethylation in 18S rRNA and 28S rRNA. Highlighted in red are the hypomethylated sites that (6 sites in 18S rRNA and 12 sites in 28S rRNA) reduce as ESCs differentiate into NSCs and neurons

| | | ESC WT | NSC WT | Neuron WT | ESC vs NSC | ESC vs Neuron | NSC vs Neuron |
|---|---|---|---|---|---|---|---|
| 18S rRNA | 116 | 0.886409 | 0.913716 | 0.972753 | ns | **** | *** |
| | 172 | 0.773057 | 0.986251 | 0.986145 | **** | **** | ns |
| | 174 | 0.62519 | 0.891424 | 0.957827 | *** | *** | ns |
| | 428 | 0.338442 | 0.762446 | 0.959955 | **** | **** | **** |
| | 468 | 0.909771 | 0.994058 | 0.997805 | ** | ** | ns |
| | 576 | 0.881844 | 0.758204 | 0.896789 | ns | ns | * |
| | 627 | 0.885702 | 0.984896 | 0.993859 | **** | **** | ns |
| | 867 | 0.723714 | 0.978765 | 0.994054 | **** | **** | ns |
| | 1272 | 0.450539 | 0.40198 | 0.833173 | ns | **** | **** |
| 28s rRNA | 1316 | 0.825662 | 0.747368 | 0.964756 | ns | ** | **** |
| | 1522 | 0.663124 | 0.994978 | 0.991648 | *** | *** | ns |
| | 1760 | 0.898756 | 0.996909 | 0.997488 | **** | **** | ns |
| | 1881 | 0.515724 | 0.603932 | 0.933649 | ns | **** | **** |
| | 2365 | 0.871694 | 0.961157 | 0.975329 | ** | ** | ns |
| | 2415 | 0.791581 | 0.918155 | 0.952379 | **** | **** | *** |
| | 2787 | 0.886671 | 0.858685 | 0.894799 | ns | ns | ns |
| | 2824 | 0.545519 | 0.986259 | 0.991524 | *** | *** | ns |
| | 2861 | 0.903707 | 0.910871 | 0.946508 | ns | ** | ** |
| | 3701 | 0.895551 | 0.993059 | 0.990555 | **** | **** | ns |
| | 3744 | 0.899003 | 0.810367 | 0.881939 | * | ns | ns |
| | 3867 | 0.398386 | 0.863865 | 0.954061 | **** | **** | ns |
| | 3944 | 0.43542 | 0.957662 | 0.971155 | **** | **** | ns |
| | 4042 | 0.827201 | 0.940962 | 0.991644 | **** | **** | *** |
| | 4306 | 0.604514 | 0.993796 | 0.965317 | **** | **** | ns |
| | 4571 | 0.872787 | 0.868315 | 0.963485 | ns | **** | **** |
| | 4590 | 0.901049 | 0.968143 | 0.997735 | **** | **** | ** |
| | 4618 | 0.593665 | 0.855277 | 0.979272 | ** | *** | ns |

**Table 2**: Sites indicating significant shift from Hypermethylation to hypomethylation in 18S rRNA and 28S rRNA

| | | ESC WT | NSC WT | Neuron WT | ESC vs NSC | ESC vs Neuron | NSC vs Neuron |
|---|---|---|---|---|---|---|---|
| 18S rRNA | 1328 | 0.974711 | 0.886104 | 0.75333 | ns | * | ns |
| | 1391 | 0.939679 | 0.991155 | 0.968983 | ** | ns | ns |
| | 1536 | 0.938834 | 0.225773 | 0.254877 | **** | **** | ns |
| 28S rRNA | 3841 | 0.995722 | 0.990531 | 0.98835 | ns | * | ns |
| | 4499 | 0.99781 | 0.993002 | 0.907781 | ns | * | * |
| | 4523 | 0.997831 | 0.982592 | 0.975999 | ns | * | ns |
| | 4620 | 0.991591 | 0.736001 | 0.852558 | **** | ** | ** |
| | 4367 | 0.874646 | 0.523633 | 0.803216 | **** | ns | **** |

**Table 3**: Sites indicating significant difference in 2’O-methylation between WT and FMR1 KO ESC 18S rRNA and 28S rRNA

| | | ESC WT | ESC FMR1 KO | P-Value |
|---|---|---|---|---|
| 18S rRNA | 116 | 0.8864 | 0.9655 | 0.00017305 |
| | 159 | 0.9833 | 0.9947 | 0.0458963 |
| | 172 | 0.7731 | 0.972 | 9.2952E-05 |
| | 174 | 0.6252 | 0.8848 | 0.0172618 |
| | 428 | 0.3384 | 0.5689 | 0.00790355 |
| | 468 | 0.9098 | 0.9919 | 0.03013167 |
| | 484 | 0.9223 | 0.981 | 0.01861479 |
| | 509 | 0.9679 | 0.9851 | 0.04543201 |
| | 576 | 0.8818 | 0.9908 | 0.02228417 |
| | 627 | 0.8857 | 0.9907 | 0.00191851 |
| | 644 | 0.9859 | 0.9956 | 0.00714317 |
| | 668 | 0.9025 | 0.9606 | 0.04182707 |
| | 867 | 0.7237 | 0.9843 | 0.00078173 |
| | 1288 | 0.9545 | 0.9685 | 0.01610787 |
| | 1328 | 0.9747 | 0.8895 | 0.00322142 |
| | 1383 | 0.9943 | 0.998 | 0.00902449 |
| | 1391 | 0.9981 | 0.9397 | 0.01181839 |
| | 1442 | 0.9353 | 0.9175 | 0.00311862 |
| | 1490 | 0.9992 | 0.9981 | 0.01562398 |
| | 1536 | 0.9388 | 0.1264 | 0.00028686 |
| | 1804 | 0.9155 | 0.9884 | 8.6571E-05 |
| 28S rRNA | 398 | 0.9978 | 0.9777 | 0.00698483 |
| | 400 | 0.9954 | 0.9853 | 0.00050524 |
| | 1326 | 0.9729 | 0.9857 | 0.04449346 |
| | 1340 | 0.8739 | 0.528 | 6.4053E-05 |
| | 1522 | 0.6632 | 0.9893 | 0.01578187 |
| | 1524 | 0.9336 | 0.9973 | 0.00019075 |
| | 1625 | 0.9884 | 0.9963 | 0.02518721 |
| | 1760 | 0.8988 | 0.992 | 0.00015771 |
| | 1881 | 0.5157 | 0.6317 | 0.01323617 |
| | 2351 | 0.9601 | 0.9977 | 0.00016025 |
| | 2363 | 0.9695 | 0.9973 | 0.00179803 |
| | 2364 | 0.9894 | 0.9963 | 0.00201424 |
| | 2422 | 0.912 | 0.9623 | 0.00548524 |
| | 2424 | 0.9685 | 0.9827 | 0.00087689 |
| | 2815 | 0.979 | 0.9907 | 0.04780965 |
| | 2824 | 0.5457 | 0.9483 | 0.00912978 |
| | 2837 | 0.9848 | 0.997 | 0.00727384 |
| | 2861 | 0.9038 | 0.9447 | 0.00043402 |
| | 3701 | 0.8956 | 0.9717 | 0.00195189 |
| | 3724 | 0.9916 | 0.9753 | 0.04275199 |
| | 3744 | 0.8988 | 0.7983 | 0.00356005 |
| | 3808 | 0.9316 | 0.9833 | 0.02257632 |
| | 3830 | 0.9567 | 0.9797 | 0.01623957 |
| | 3867 | 0.3985 | 0.8123 | 0.00467805 |
| | 3869 | 0.9173 | 0.778 | 0.03731292 |

| | | | | |
|---|---|---|---|---|
| | 3887 | 0.9781 | 0.9993 | 0.02039052 |
| | 3925 | 0.9369 | 0.9753 | 0.01006758 |
| | 3944 | 0.4353 | 0.9223 | 0.0010103 |
| | 4042 | 0.8273 | 0.9603 | 0.00028329 |
| | 4227 | 0.9256 | 0.9933 | 0.00339875 |
| | 4228 | 0.9638 | 0.9967 | 0.00122037 |
| | 4306 | 0.6046 | 0.8987 | 8.2345E-05 |
| | 4456 | 0.9934 | 0.9637 | 0.0022907 |
| | 4499 | 0.9979 | 0.9377 | 0.03272395 |
| | 4523 | 0.9978 | 0.9903 | 0.00519714 |
| | 4571 | 0.8729 | 0.936 | 0.03591427 |
| | 4590 | 0.901 | 0.9483 | 0.00595971 |
| | 4618 | 0.5938 | 0.9 | 0.01413858 |
| | 4620 | 0.9915 | 0.8897 | 0.00535085 |
| | 4637 | 0.8744 | 0.5803 | 0.00145973 |

Table 4: Sites indicating significant difference in 2'O-Methylation between WT and FMR1 KO NSC 18S rRNA and 28S rRNA

| | | NSC WT | NSC FMR1 KO | P-Value |
|---|---|---|---|---|
| 18S rRNA | 99 | 0.9947 | 0.9967 | 0.01071522 |
| | 116 | 0.9137 | 0.96 | 0.0007347 |
| | 159 | 0.9754 | 0.9814 | 0.02030539 |
| | 166 | 0.9902 | 0.9943 | 0.04352719 |
| | 172 | 0.9863 | 0.99 | 0.02795788 |
| | 428 | 0.7624 | 0.8545 | 0.00086674 |
| | 512 | 0.9785 | 0.9842 | 0.01070892 |
| 28S rRNA | 398 | 0.9977 | 0.9949 | 0.00781102 |
| | 1326 | 0.9886 | 0.9921 | 0.0102442 |
| | 2415 | 0.9182 | 0.8871 | 0.00247807 |
| | 2861 | 0.9109 | 0.8917 | 0.00568395 |
| | 3760 | 1 | 0.9976 | 0.03563853 |
| | 4618 | 0.8553 | 0.8221 | 0.01716329 |

Table 5: Sites indicating significant difference in 2'O-Methylation between WT and FMR1 KO neuron 18S rRNA and 28S rRNA

| | | Neuron WT | Neuron FMR1 KO | P-Value |
|---|---|---|---|---|
| 18S rRNA | 99 | 0.9925 | 0.9978 | 0.00376031 |
| | 172 | 0.9858 | 0.995 | 0.00212042 |
| | 436 | 0.9385 | 0.9685 | 0.04156871 |

| | | | | |
|---|---|---|---|---|
| | 627 | 0.9938 | 0.9973 | 0.02739604 |
| | 799 | 0.974 | 0.9953 | 0.00414657 |
| | 1804 | 0.9965 | 0.9983 | 0.04479426 |
| 28S rRNA | 1871 | 0.9993 | 0.9968 | 0.03076601 |
| | 1881 | 0.9338 | 0.8988 | 0.0017439 |
| | 2861 | 0.9463 | 0.9235 | 0.04291374 |
| | 3899 | 0.991 | 0.9955 | 0.00692005 |
| | 4054 | 0.989 | 0.9773 | 0.03487105 |
